# Reading positional codes with fMRI: Problems and solutions

**DOI:** 10.1101/076554

**Authors:** Kristjan Kalm, Dennis Norris

## Abstract

Neural mechanisms which bind items into sequences have been investigated in a large body of research in animal neurophysiology and human neuroimaging. However, a major problem in interpreting this data arises from a fact that several unrelated processes, such as memory load, sensory adaptation, and reward expectation, also change in a consistent manner as the sequence unfolds. In this paper we show that the problem of extracting neural data about the structure of a sequence is especially acute for fMRI, which is almost exclusively the modality used in human experiments. We show that such fMRI results must be treated with caution and in many cases the assumed neural representation might actually reflect unrelated processes.

## 1 Introduction

One of the most important features of human cognition is the ability to bind individual events into a sequence. Almost any complex task requires us to remember not only the individual elements but also the order in which they occurred. For example, two tasks such as starting a car and stopping it might share the same events but in different order. All computational models of sequence processing acknowledge this distinction between the representations of items in memory and the representation of the order in which they occur (Henson & Burgess, 1997; Page & Norris, 1998). The view that item’s position in the sequence is encoded separately and independently of their identity has been also suggested by decades of research in human behaviour and animal neurophysiology.

Neurons in the monkey prefrontal cortex (PFC) have been found to be selective for each position in a learned sequence (Nakajima, Hosaka, Mushiake, & Tanji, 2009; Averbeck, Crowe, Chafee, & Georgopoulos, 2003; Inoue & Mikami, 2006; Naya & Suzuki, 2011). Figure 1C gives an example of a simple positional code showing the responses of position-sensitive neurons from monkey supplementary motor area as observed by Berdyyeva and Olson (2010). Other research on animal neurophysiology has suggested that the hippocampus encodes the position of items in a sequence (Mankin et al., 2012; Manns, Howard, & Eichenbaum, 2007a; Naya & Suzuki, 2011; Pastalkova, Itskov, Amarasingham, & Buzs´aki, 2008), with some authors proposing the existence of ’time cells’ tracking the temporal position of items in a sequence (MacDonald, Lepage, Eden, & Eichenbaum, 2011; MacDonald, Carrow, Place, & Eichenbaum, 2013). From hereon we refer to such neural representation of the item’s position in the sequence as *positional code*. The extensive literature on the neural representation of the positional code is summarised in Table 1.

However, interpreting a neural signal tracking the positional code suffers from a major methodological problem: items in different positions necessarily differ along other dimensions too. For example, in a memory task, memory load will be greater at position three than position two. Changes in neural activity that are sensitive to memory load might therefore give the appearance of coding position. An item in position *n* will always be associated with a load of *n* items. Any neural index of load will therefore consistently be in a different state for items in different positions. An item in position *n* also occurs at a later time than item *n* − 1. Sensory adaptation might change the neural response to items as the sequence progresses. Such a signal could also masquerade as a positional code. Any or all of these factors might therefore lead to a differential neural response which would correlate with the position of an item in a sequence, but which might play no role in determining how the brain codes temporal position. In their analysis of how we can measure information in the brain, De-Wit, Alexander, Ekroll, and Wagemans (2016) made a contrast between ”cortex as receiver” and ”experimenter as receiver”. There may be ways in which we as experimenters can decode neural states to recover information about temporal position, but what we would like to do is to identify specifically those neural representations that the cortex uses to represent temporal position and to drive behaviour.

**Figure 1:**
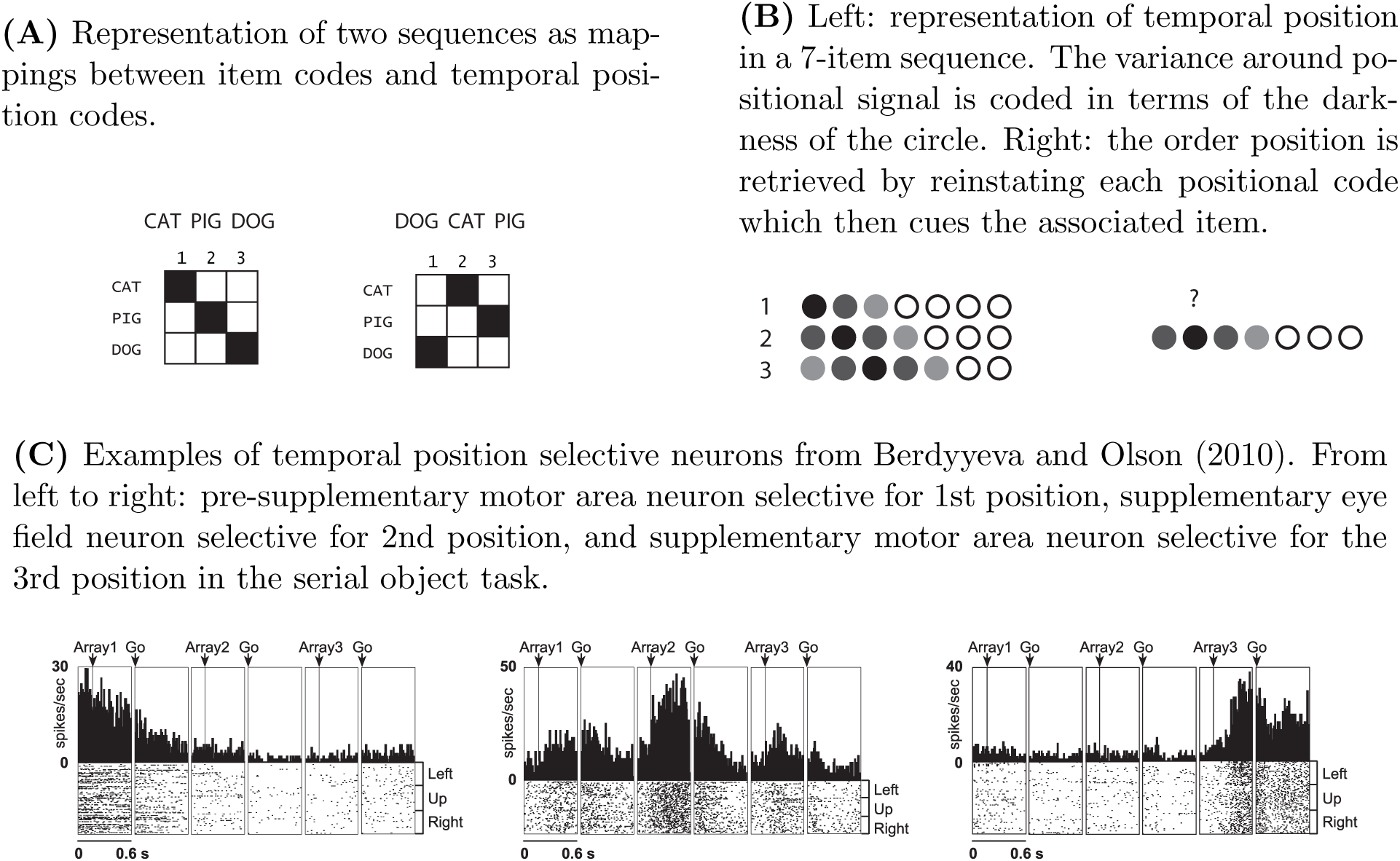
Sequence representation and temporal position

In this paper we show that dissociating positional ’read-out’ from a neural positional code is an especially difficult problem for fMRI data. We show that fMRI data acquired from sequentially presented stimuli suffer from several confounds. First, we show that with any sequence processing task there are experimental variables which are collinear with the positional signal (e.g. memory load, sensory adaptation, etc.) and which can serve as a positional code. Second, we show how interference between stimulus representations, task phases, and measurement modalities can also lead to a similar positional read-outs indistinguishable from a dedicated positional code. Importantly these correlated effects do not simply result in a univariate change in signal that varies across sequence position but also change the pattern of information that can be read out by multivariate methods.

The problem of interpreting a positional signal is especially relevant since neural data on human sequence processing comes almost exclusively from fMRI studies (Table 1). Our simulations and experimental data show that results from fMRI experiments studying the positional code must be treated with caution. Specifically, in many cases the assumed positional code might actually reflect processes which are correlated with position in the sequence instead.

## 2 Positional code from collinear processes

Any signal tracking the position of an item in a sequence will be collinear with a number of cognitive processes:

- Memory load – signal for position *n* will always co-occur with a memory load of *n* items when storing a sequence. Any neural index of load will therefore always reflect the progression of sequence.
- Sensory adaptation – neural responses in the human sensory cortex have been shown to monotonically decrease as a response to sequentially presented stimuli (Henson & Rugg, 2003; Summerfield, Trittschuh, Monti, Mesulam, & Egner, 2008; Larsson & Smith, 2012). Any signal that monotonically changes over sequence positions can be used to read out position-like code.
- Reward – in most animal studies the subject is rewarded after successfully attending or recalling a sequence. This means that the next item in a sequence is always closer to the reward. Neurons tracking the temporal proximity of reward have been described in both monkey and rodent studies (Berdyyeva & Olson, 2011; MacDonald et al., 2011; Naya & Suzuki, 2011).
- Passage of time – signal for position *n* always occurs after the signal for position *n* − 1.

All these processes represent a change in the cognitive state of the participant throughout the processed sequence, and hence will necessarily be collinear with any positional code. It follows that in the analysis of experiments on temporal order it is necessary to distinguish between a dedicated positional code and a positional read-out from collinear processes.

Next, we provide two examples of positional read-out based on human fMRI data. In the first example we show how sensory adaptation in the sensory cortices can be interpreted as a positional signal. In the second example we show how differences in retinotopic activation over the course of the sequence can similarly provide positional read-out. In the final part of the section we provide simulations which explore whether it is possible to develop methods to subtract the effects of such collinear processes from sequentially obtained data.

### 2.1 Sensory adaptation

Sensory adaptation across sequence positions has been observed in a number of fMRI studies of sequence processing as a decreasing univariate signal over positions (Summerfield et al., 2008; Larsson & Smith, 2012; Henson & Rugg, 2003). Note that an inverse trend, where the univariate signal increases over sequence positions, has also been observed (Kalm & Norris, 2016). The latter most likely reflects the attenuation of the BOLD signal in response to sequentially presented stimuli as reported in other fMRI studies on human STM (Rottschy et al., 2012; Wager & Smith, 2003). However, the direction of the univariate change is unimportant as any consistent change over sequence positions will permit position decoding.

Here we used two human fMRI datasets obtained with a sequence processing task (Kalm & Norris, 2014, 2016) to carry out a classification analysis of item position in a sequence. In both cases we chose the sensory cortex of the presented stimuli as a region of interest (ROI): in the first experiment the sequences were presented auditorily (Kalm & Norris, 2014) and in the second visually (Kalm & Norris, 2016). Since in both experiments sensory areas responded differentially to sequence positions (Figure 2A) linear classification analysis can be used to predict the position of the items significantly above chance (Figure 2B). However, in both cases the signal changes were uniform across all voxels in the anatomical region suggesting not a dedicated positional code, but sensory adaptation or change in measurement noise. Sensory adaptation thus serves as a clear example how a monotonically changing signal can be read out by an experimenter as a positional code.

**Figure 2:**
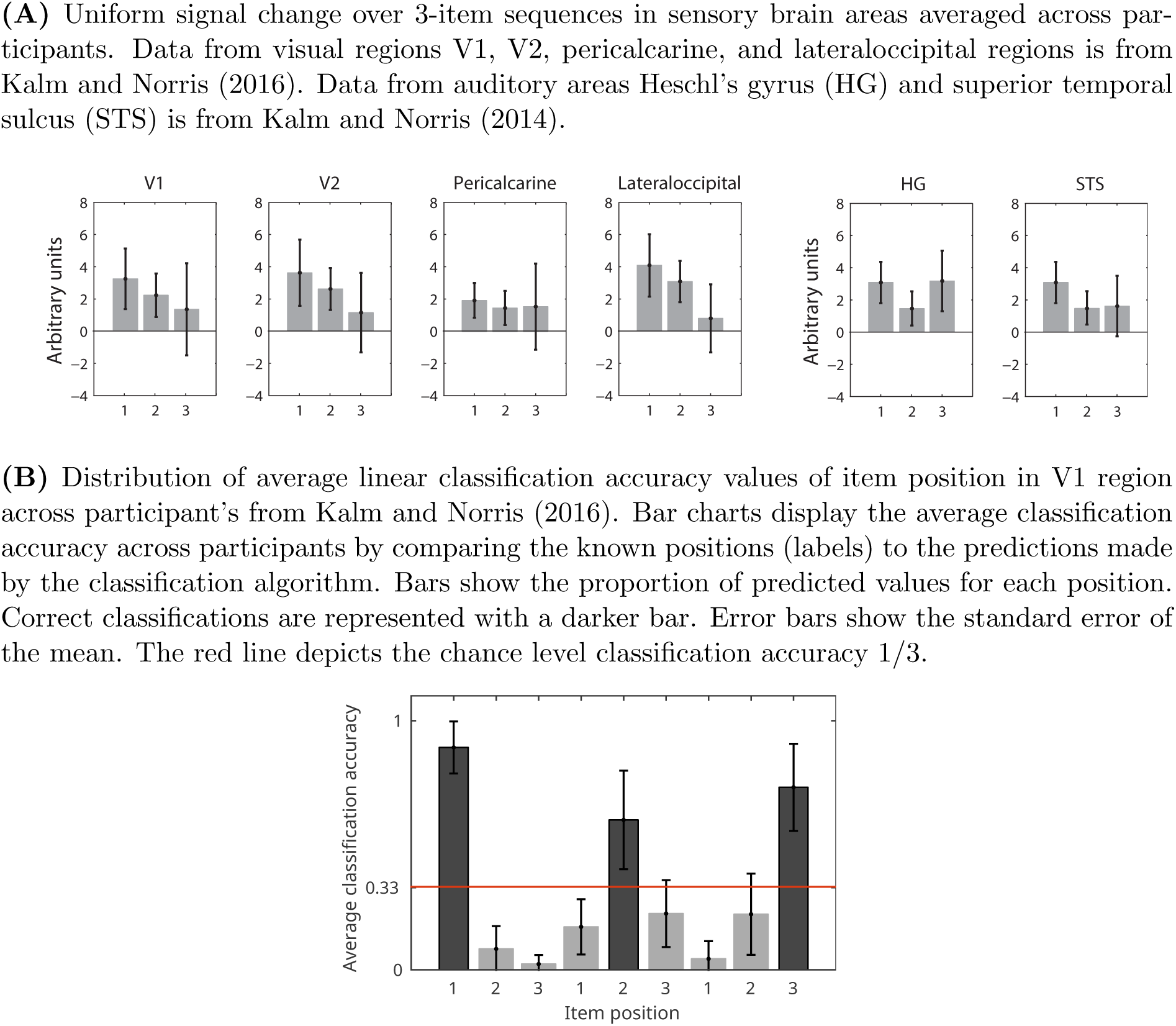
Sensory adaptation in the sensory cortex and decoding order position.

### 2.2 Retinotopic activation

In the example above (*Sensory adaptation*) the population of neural units (sensory cortex) responded uniformly to sequentially presented stimuli. Next we present a case where neural units *within* the population respond differentially across the sequence. We use fMRI data from a visual sequence processing task to show that the response in the primary visual cortex can be used to predict the position of the item in the sequence. However, this is possible not because of any positional code but because of task-selective voxels in the visual cortex.

In Kalm and Norris (2016) participants had to attend a sequence of visually presented images followed by a manual response indicating the order of the items (Figure 16). Importantly, all images were controlled for luminance and cropped to ensure that each image appeared in a similar retinal area: all stimuli subtended a 6° visual angle around the fixation point in order to elicit an approximately foveal retinotopic representation. As a result, all sequence items elicited approximately similar retinotopic response in the foveal area of the primary visual cortex.

The authors observed that the activation of the retinotopically driven voxels was correlated with the relative suppression of the voxels outside of the retinotopically activated areas (Figure 3A). Such suppression has been observed as a function of stimulus location in the visual field (Maier et al., 2008) and attention (Gouws et al., 2014; Smith, Singh, & Greenlee, 2000). Importantly, the amount of activation and suppression changed across sequence positions. Since the sequence items were presented in immediate succession, the extent of retinotopic suppression and activation varied as a function of item’s position in the sequence. Figure 3B shows data from a single participant’s V1, where voxels are split into two groups: retinotopically activated (red-yellow on Figure 3A) and suppressed (blue-cyan on Figure 3A) represented by red and blue lines. As the activation and suppression of two different sets of voxels changes across positions, a linear classification algorithm can use the difference between activated and suppressed voxels, or the difference between the red and blue lines on Figure 3B, to reliably predict the item’s position.

This can be further illustrated when linear discriminant analysis (LDA) class boundaries based on item position are plotted with following sets of voxels from V1:

1. All voxels (including both retinotopically activated and suppressed voxels)
2. Only activated voxels (*p* < 0.01)
3. Only suppressed voxels (*p* < 0.01)

LDA shows that the linear classifier is only able to reliably predict the position of the item when both activated and suppressed voxels in the brain region are included (Figure 4, top row). The classification is at chance level if only one set of voxels are used (Figure 4, row 2-3).

**Figure 3:**
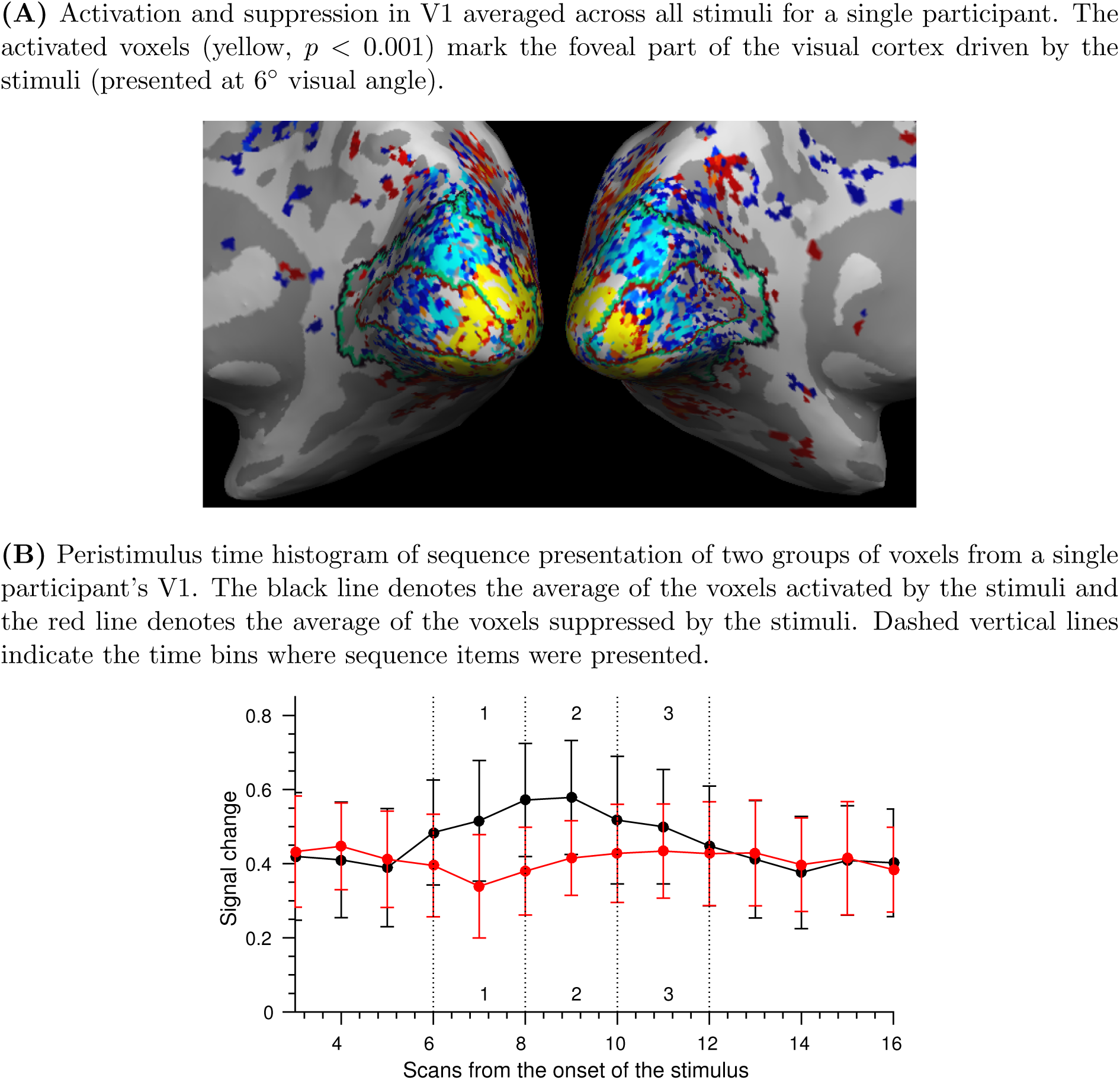
Interference between task phases: retinotopic suppression

Next we carry out a simulation of sequentially generated fMRI data to explore whether both uniformly and differentially proceeding collinear processes could be controlled for when trying to extract a positional code.

**Figure 4:**
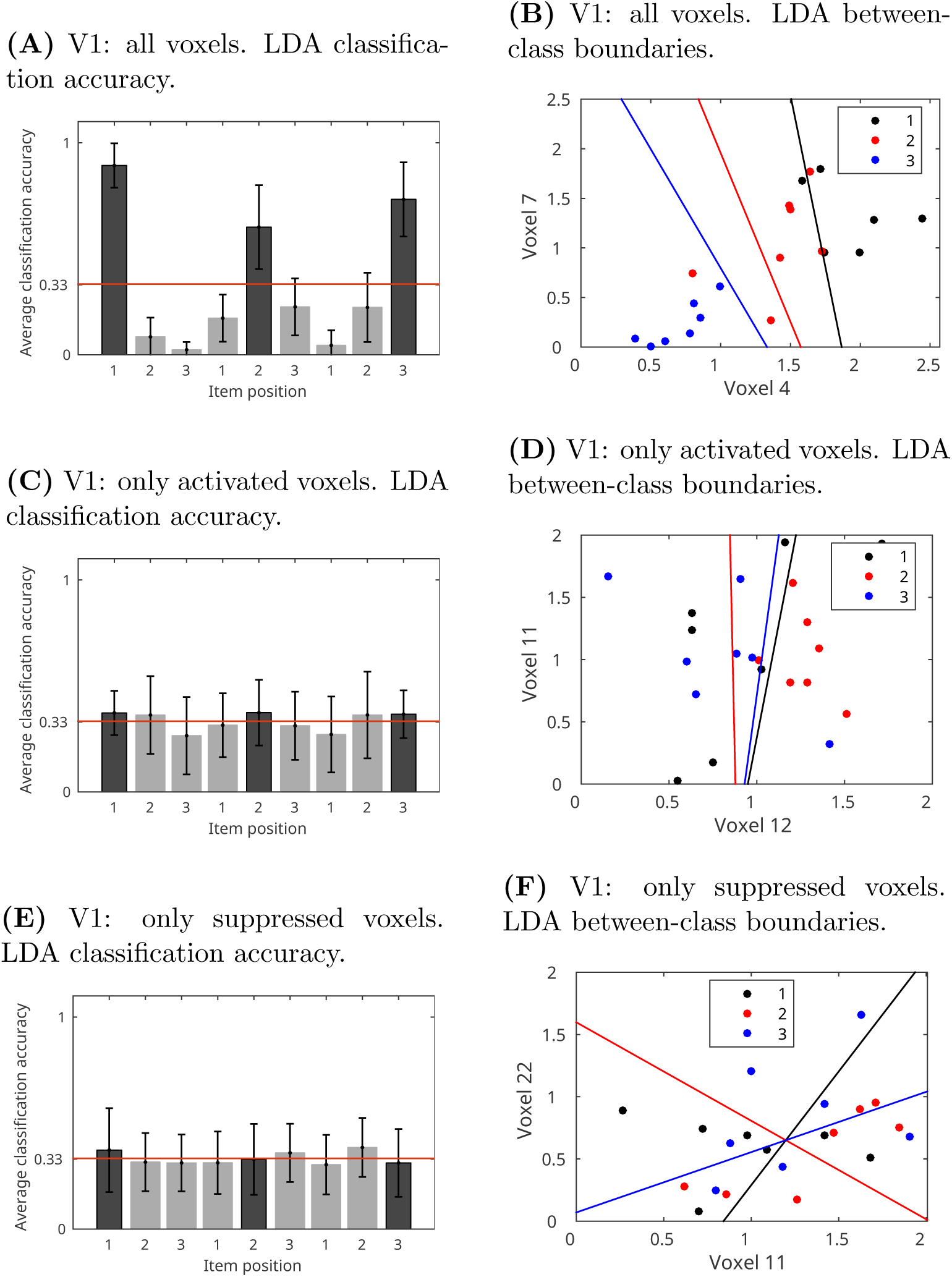
LDA of item position in V1 using different subsets of voxels. *Top row*: all voxels from V1; *middle row*: only retinotopically activated voxels from V1; bottom row: only retinotopically suppressed voxels from V1. *Left column*: Bar charts display the average classification accuracy across participants by comparing the known positions (labels) to the predictions made by the classification algorithm. Bars show the proportion of predicted values for each position. Correct classifications are represented with a darker bar. Error bars show the standard error of the mean. The red line depicts the chance level classification accuracy 1/3. *Right column*: LDA between-class boundaries based on two voxels from the set. Data from Kalm & Norris (2016).

## 3 Simulation of collinear processes

Here we simulate two types of position-collinear processes which can serve as a positional readout. In the first case the brain area responds uniformly along the sequence (e.g. sensory adaptation) and in the second case units within the population respond differentially. We show that in the first case we can make reasonable *a priori* assumptions about the nature of the positional code and hence remove a uniform signal. However, when the population responds differentially to sequence positions there are no prior criteria to distinguish positional read-out from a positional code.

The MATLAB/Octave code for the simulated data and plots is freely available at http://imaging.mrc-cbu.cam.ac.uk/imaging/KristjanKalm/poscode/.

### 3.1 Uniformly changing signal across sequence positions

Here we model sensory adaptation in a simple sequence processing task as an example of a uniformly changing position-collinear process. We show how human fMRI data obtained with the same task fits the simulation results. We also propose a data pre-processing step – demeaning of neural responses – as a tool to eliminate univariate signal collinear to the positional code.

Throughout the simulations we use the term ’brain region’ for a population of neural units and the term ’voxels’ for units themselves. This makes the terminology compatible with the experimental data presented from human fMRI experiments.

#### 3.1.1 Representation of sequence items in a brain region

As a baseline condition we simulate the case where the only information stored in a brain region is item information (without any positional code) and where there is no position-collinear information such as decay or interference. We simulate a sensory brain region of *n* = 20 voxels which encodes identities for three different items *i* as independent samples from the uniform distribution (Figure 5A):

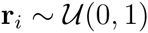

A brain region’s response Y to the item *i* will be the item pattern r_*i*_ plus some noise sampled from *n*-dimensional Gaussian distribution with a zero mean.

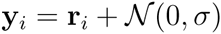

To model a noisy average of these patterns we simulate an experiment where those three items are presented in different order as sequences for 6 times. The simulated response matrix Y depicts those 6 sequences with item and position values labelled on the x-axis (Figure 5B).

**Figure 5:**
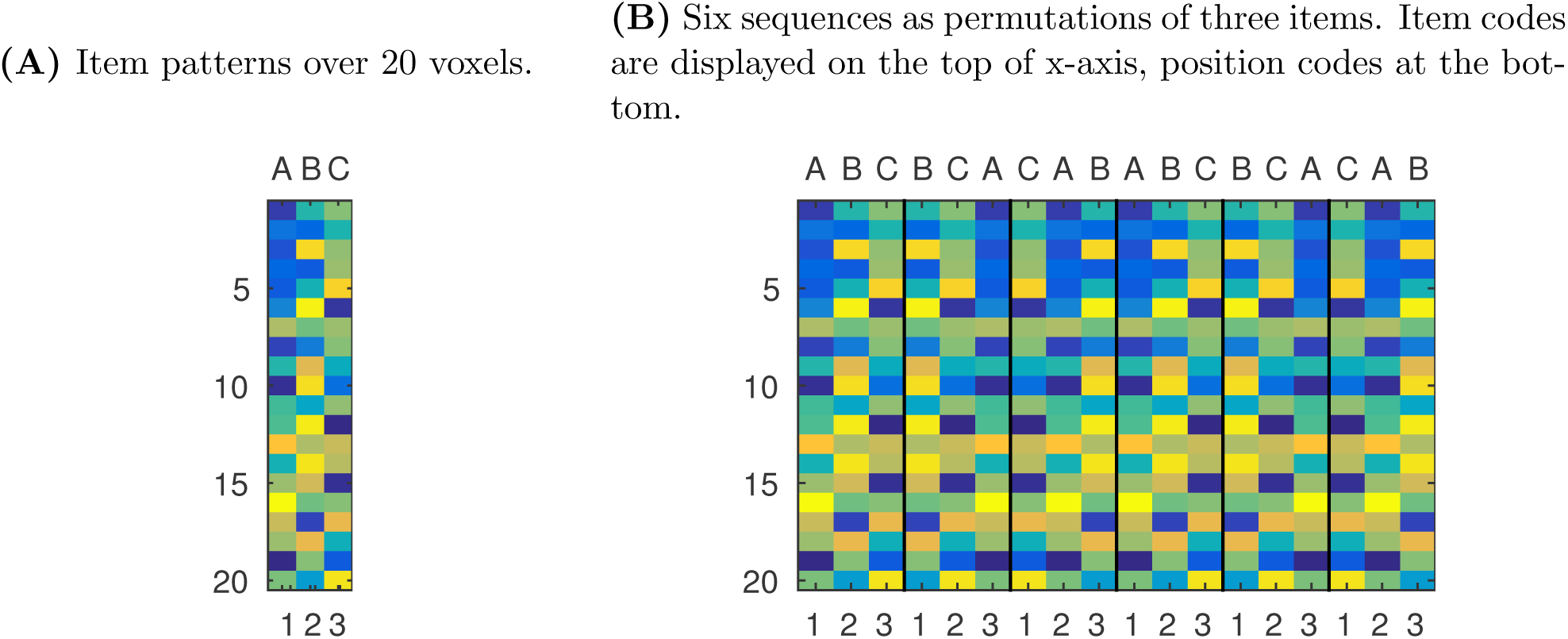
Simulated responses to items

As a result, the brain region’s response matrix Y contains noisy representations of item identity but no information about position in the sequence. This can be visualised by plotting the scatter of the data Y and LDA class borders according to item and position labels (Figure 6). It is obvious that patterns Y are only linearly separable in terms of item identity (Figure 6A) but not position (Figure 6B).

**Figure 6:**
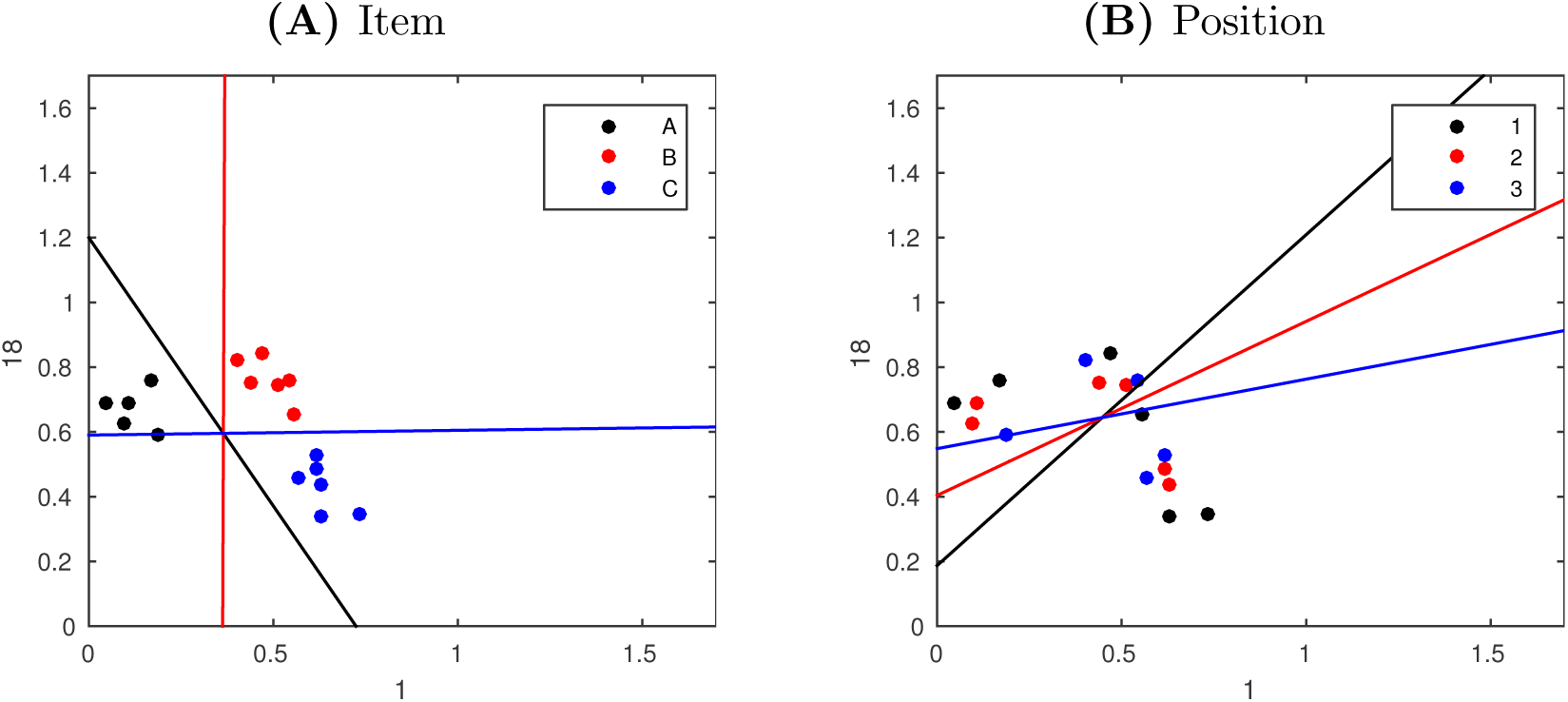
The scatter of item patterns and LDA between-class boundaries based on the two most informative voxels.

#### 3.1.2 Sensory adaptation

So far we have assumed that item representations are completely independent of sequence position. Next we consider the case where there is a degree of sensory adaptation across the sequence. We simulate sensory adaptation for a brain region as a fixed vector across voxels multiplied by a decreasing function of sequence position, plus a Gaussian noise of fixed magnitude. This means that sensory adaptation will influence all voxels in the brain region similarly. In other words, in terms of a neural response of a brain region, sensory adaptation is a univariate signal decreasing monotonically over sequence positions.

We simulate sensory adaptation for all voxels i.e. voxels respond to stimulus positions {1, 2, 3} by a decreasing vector a = [1, 0.7, 0.4]. The average responses of the voxels can be shown as column-wise means of the response matrix (Figure 7B). As a result, the response of the brain region allows us to linearly separate both item identities and their positions in the sequence (Figure 7C). Re-running this simulation 250 times yields a distribution of average LDA accuracy values (Figure 7D).

In sum, simulating sensory adaptation on top of independent item codes allows us to ’decode’ items in terms of their positions since it is predicted by the amount of sensory adaptation. The same effect would result from any cognitive process collinear to the item position such as increasing memory load or passing time.

**Figure 7:**
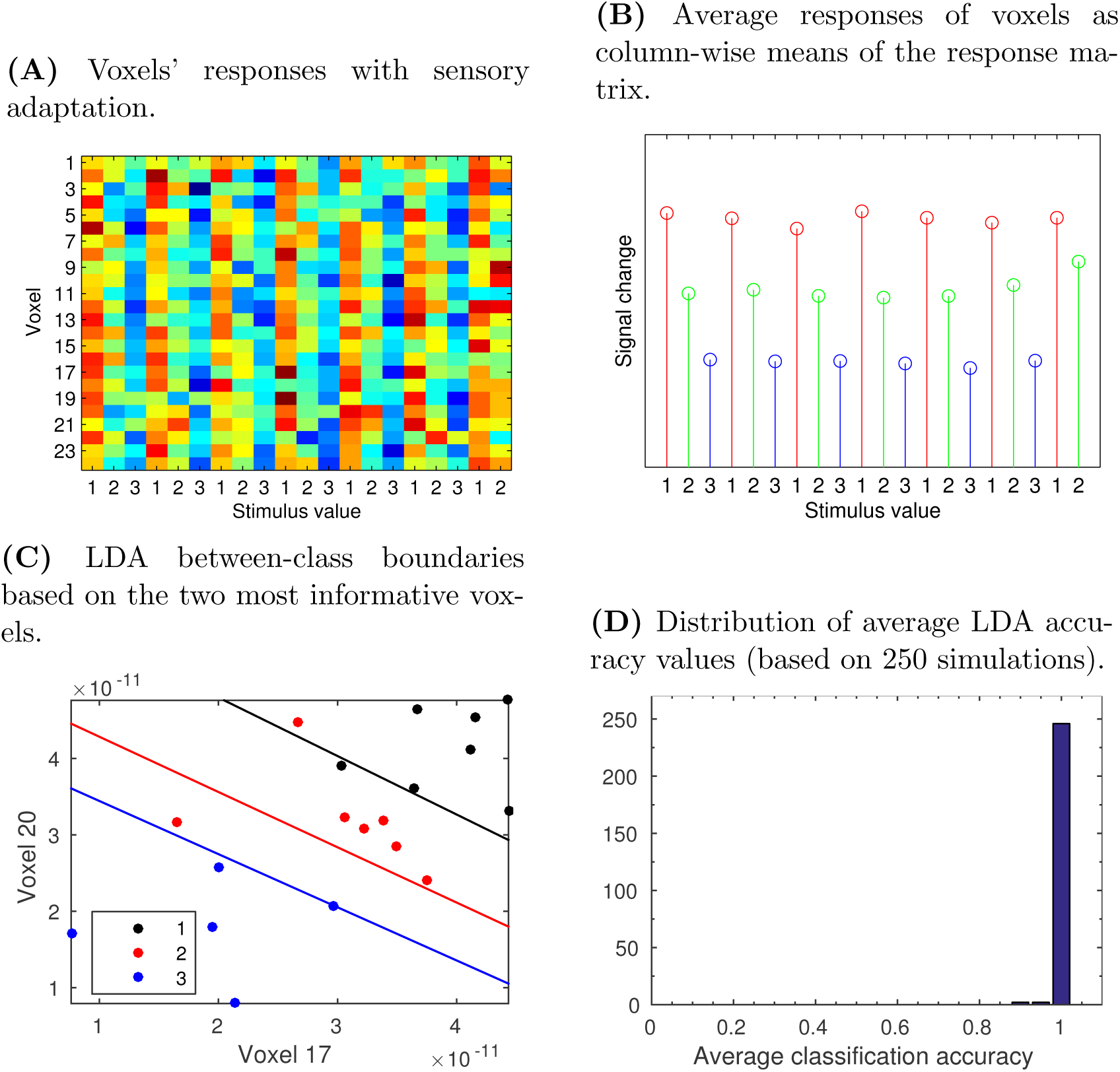
Simulation of sensory adaptation

#### 3.1.3 Eliminating uniform signal by de-meaning

If we assume that any collinear process to sequence position affects all voxels in the brain region uniformly then simple de-meaning of the response matrix will eliminate any univariate signal from the data.

Here we z-score the response matrix before classification so that column-wise averages equal zero and values of the matrix correspond to z-scores based on the column mean (Figure 8). Carrying out LDA as before shows that the resulting average classification accuracy is at chance level as z-scoring the response matrix removes effects common to all voxels. Similarly, when z-scoring was applied to the fMRI data above (see *Sensory adaptation*), positional effects were no longer significant.

**Figure 8:**
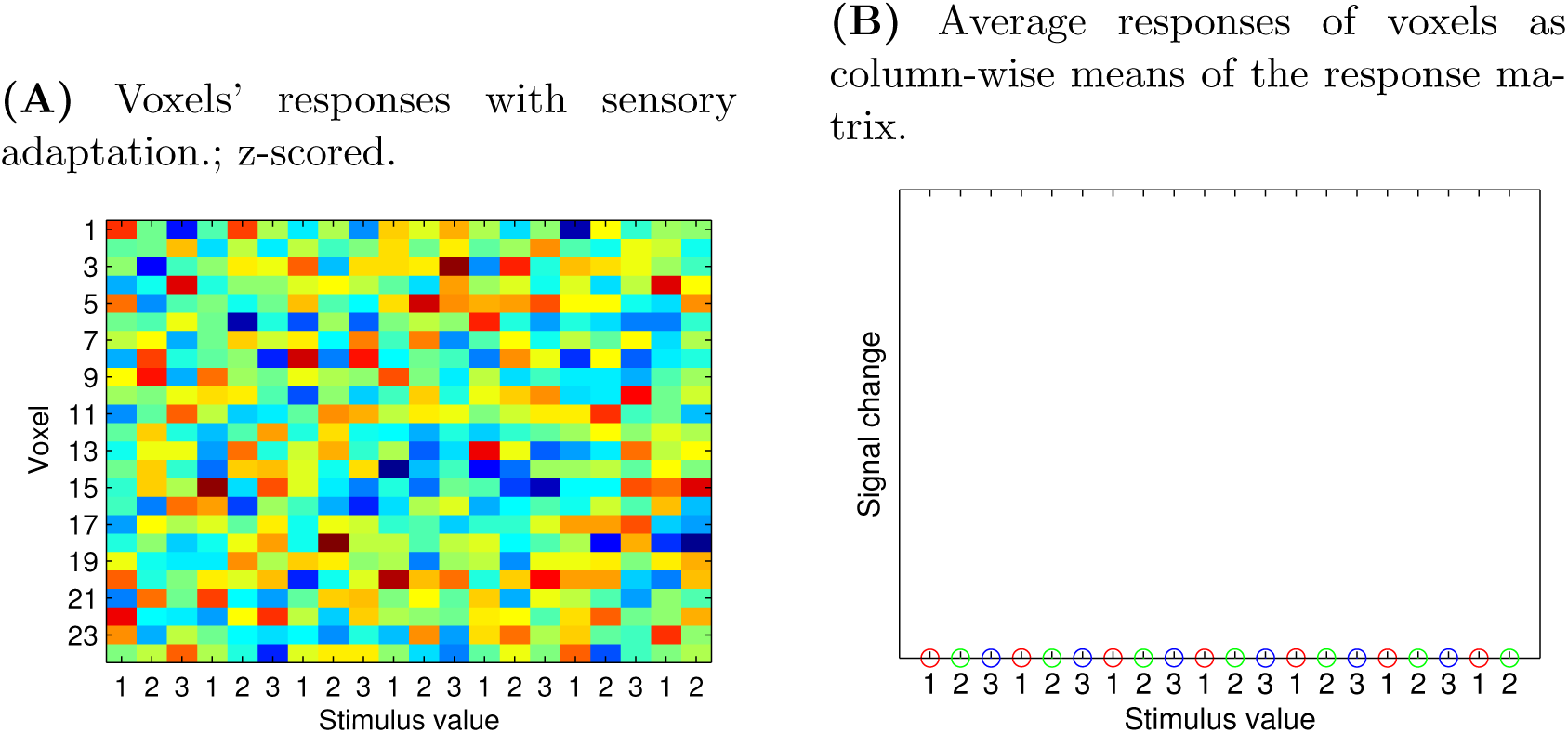
Simulation of sensory adaptation, z-scoring

However, is is also possible that the sensory brain region actually contains a positional code. It is clear that in order to survive a de-meaning process a dedicated positional code must not be uniform across voxels. De-meaning process cannot affect a multivariate positional signal which affects voxels differentially. We can model each voxel’s position preference *T* as a Gaussian likelihood function over the position values of the stimuli: i.e. each voxel responds most to a single position and less to adjacent positions: *T ∼ N* (*Position, σ*), (note that alternative tuning distributions are also feasible, see the simulation code for examples). Next, we add sensory adaptation (Figure 9B), Gaussian noise (Figure 9C), z-score the data (Figure 9D), and carry out LDA, as above. The resulting average classification accuracy will be close to 100%: since z-scoring does not affect voxel pattern similarity, the positional code is used by the linear classifier to successfully distinguish between order positions.

**Figure 9:**
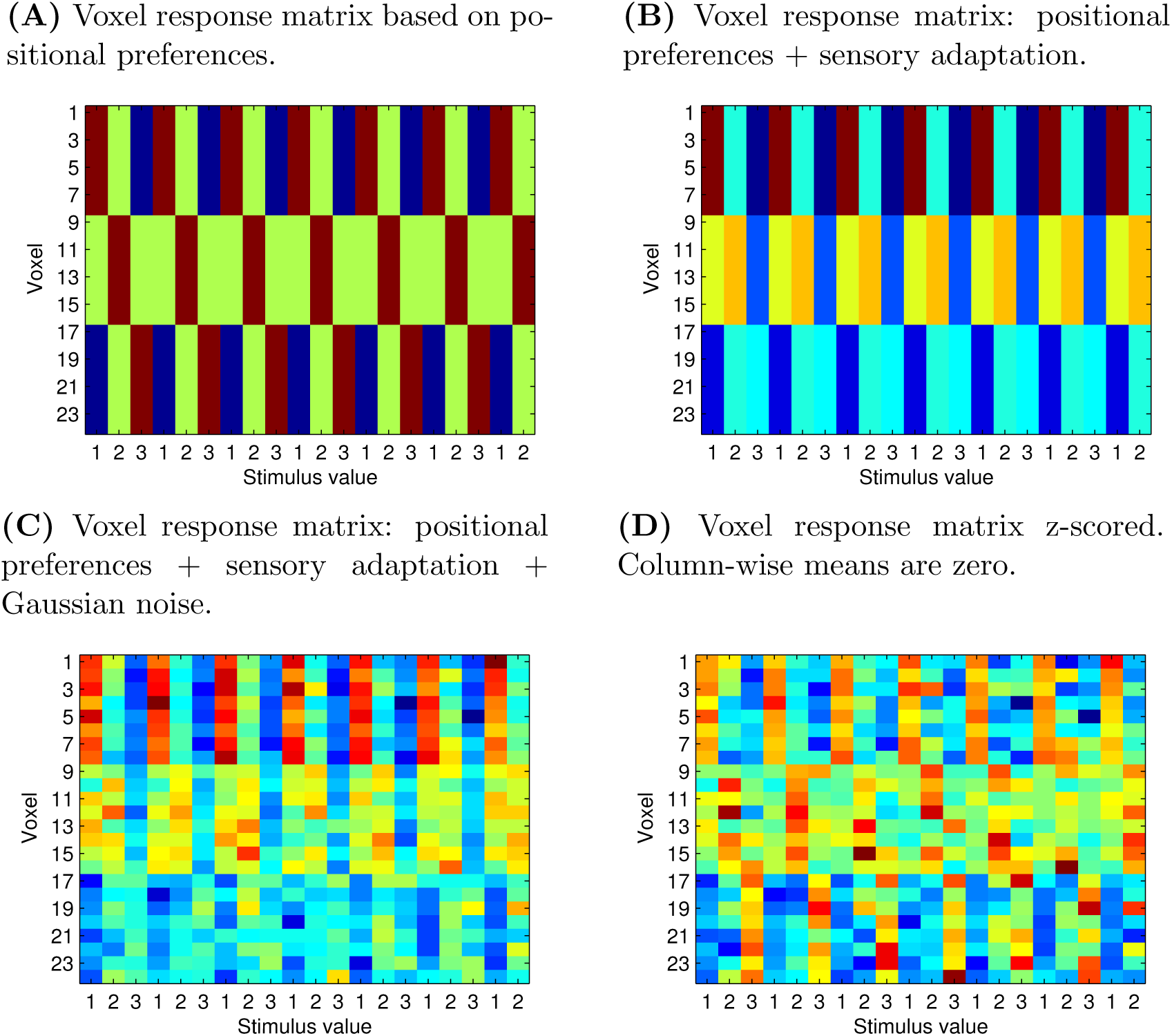
Simulation of sensory adaptation

However, any differential response within the brain region to sequence positions – such as the retinotopic activation example above – will similarly remain unaffected by z-scoring. As a result we can use de-meaning only to remove uniform effects from the brain region’s response.

### 3.2 Summary of position-collinear effects

A number of cognitive processes take place while stimuli are processed in a sequence. Importantly, several of them – time, memory load, sensory adaptation – will be collinear to any signal tracking the position of items in a sequence.

We showed that uniform position-collinear processes – such as sensory adaptation– can be subtracted from neural responses by a de-meaning technique such as z-scoring. Importantly, this relies on an assumption that such processes will influence all units uniformly in a neural population. However, if individual voxels within a brain region respond differentially – such as in the case of retinotopic activation – the neural response becomes indistinguishable from a dedicated positional code.

## 4 Positional code from interference

A positional ’read-out’ without a dedicated positional code can also arise from interference between sequentially presented stimulus representations. Here we use a simulation to show that a model of sequence representation which only includes item codes and no dedicated positional code can elicit positional effects given some interference between item codes. To illustrate this, imagine a brain region where the representations of successive items are overlayed on top of each other. Each successive item elicits a neural pattern that is a mixture of its own representation and a decaying representation of the preceding items. Such superimposed items could be linearly separable in terms of their positions alone without the need of any explicit representation of position.

Here we look at two cases of interference between item codes – additive and proportional interference– and how both can lead to position-like codes. The item representations are modelled exactly as above in *Representation of sequence items in a brain region*.

### 4.1 Additive interference

Interference between representations can occur when the state of the memory is not completely wiped clean every time a new stimulus arrives. Instead, the new state of the memory might be a mixture of the new stimulus and the previous state of the memory. Here we assume that at sequence position *p* the response of the brain region Y equals to the item pattern r_*i*_ plus some residual activity from the previous state of the brain region:

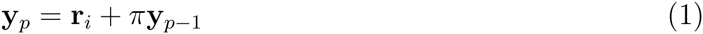

where *p* is the position of the item in the sequence and *π* is the mixing coefficient which determines the proportion of the residual activity. Here *π* declines with a constant rate over previous states of Y so that:

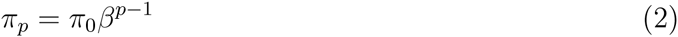

where *β* is the rate parameter of the decreasing mixing coefficient *π*, and the initial value of *π*_0_ = 1. Setting the initial value of *π* to 1 ensures that the current item pattern is always represented in full. To illustrate this mechanism consider two different *β* values and how they affect interference in a 3-item sequence ’CBA’:

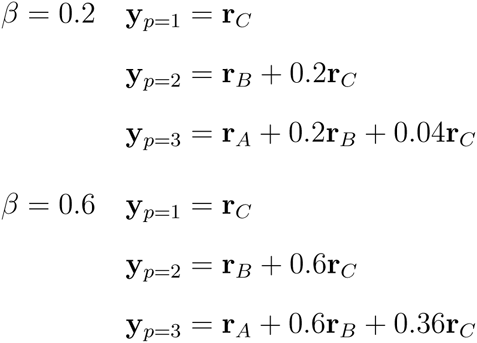

It is clear that the value of the *β* parameter determines the amount of interference from previous items: when *β* = 0 there is no interference, and when *β* > 1 the activity from previous items contributes more to the current activity pattern y_*p*_ than the current item pattern r_*i*_.

Importantly, with each arriving item the overall activity of the brain region, as defined by the vector sum of y_*p*_, increases, since some of the previous response is added to the new response. In other words, additive interference as defined above (Eq. 1) guarantees that:

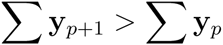

Similar increase in brain activity as a function of the number of sequentially presented items has been observed in several neuroimaging studies of short-term memory (Rottschy et al., 2012; Wager & Smith, 2003).

#### 4.1.1 Additive interference enables position decoding

If we simulate additive interference as described above then despite the brain region only encoding item identity information we can linearly separate patterns Y in terms of their position because the total activity increases as a function of position. The effect of additive residual activity on sequence positions can be shown by plotting the positional means before and after interference transform (Figure 10).

**Figure 10:**
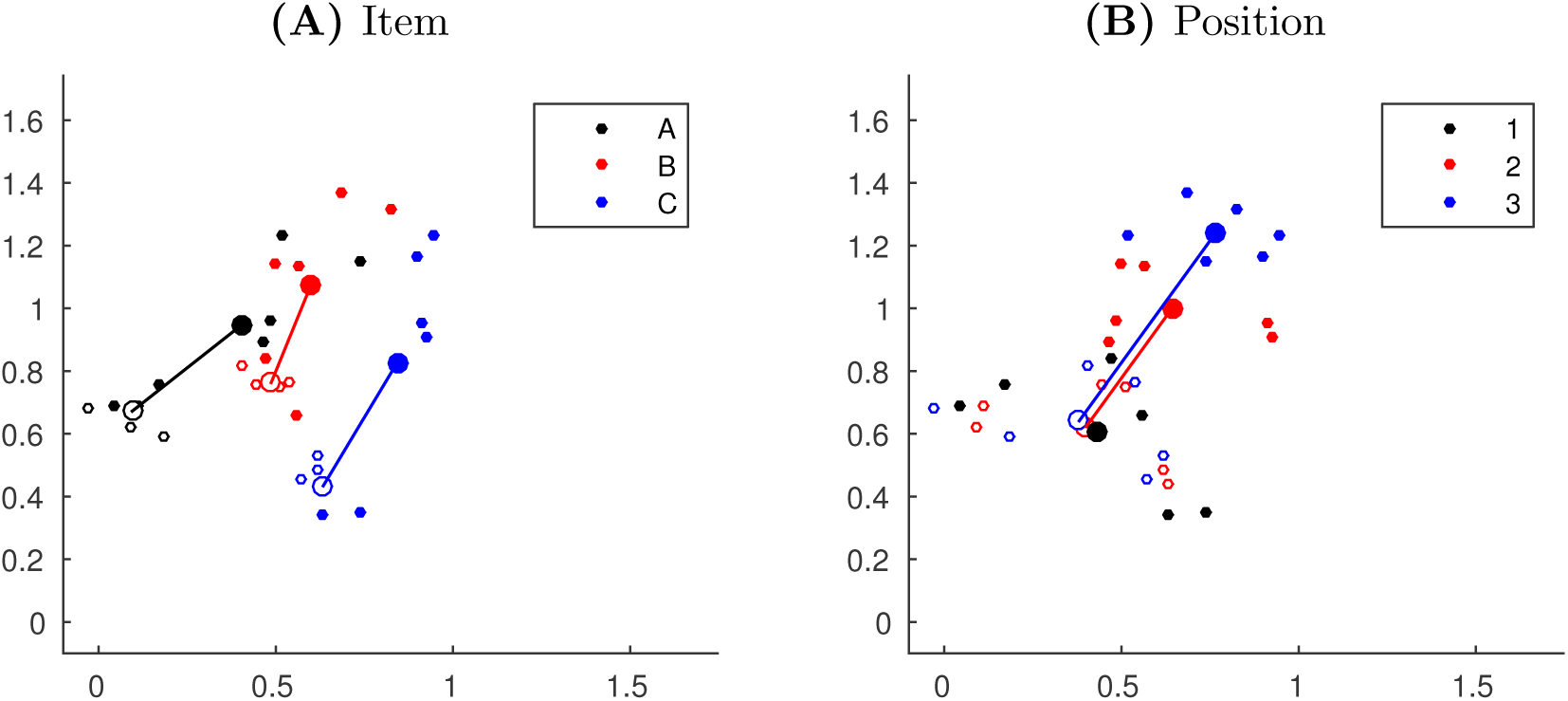
The transformation of response values for two voxels as a result of interference (*β* = 0.5). Small circular markers depict response patterns, larger circular markers depict pattern means. Empty markers depict the original patterns and means, filled markers depict the data after simulating the interference process. Solid lines depict the movement of the class means as a result of interference.

Note that patterns pertaining to the first positions in the sequence (black markers on Figure 10B) have not moved since there is no interference for the first items in the sequence from previous items. The position-wise transformation of the response patterns allows to separate them linearly using both item and position labels (Figure 11).

**Figure 11:**
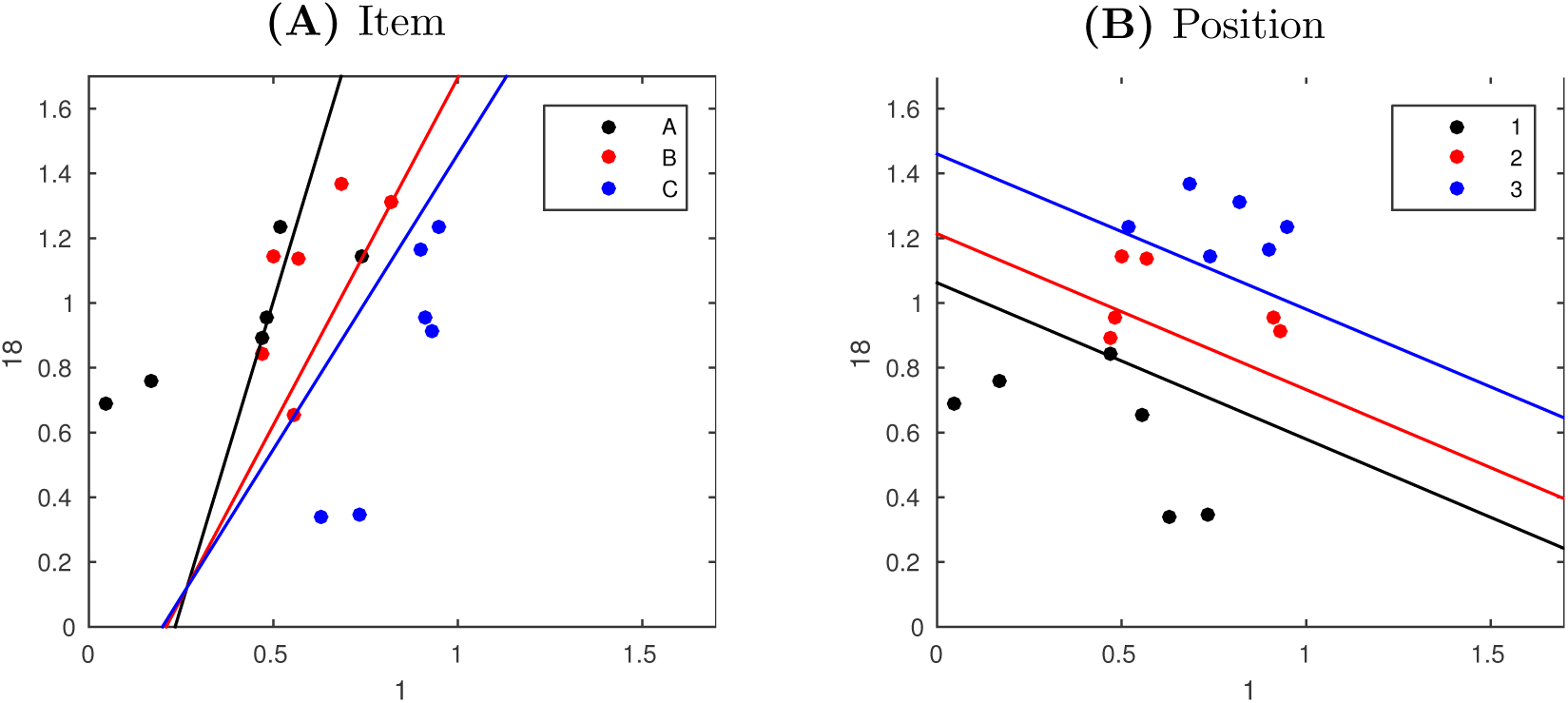
LDA between-class boundaries for two voxels, interference *β* = 0.5

We can now decode the position of the items significantly above chance because item position correlates with the amount of response in the simulated brain region. Plotting the classification accuracy of both item and position as a function of interference (*β* parameter value, Eq. 2) we can see that even with relatively small *β* values positional decoding becomes significantly greater than chance whilst it is always possible to decode item identity above chance (Figure 12).

**Figure 12:**
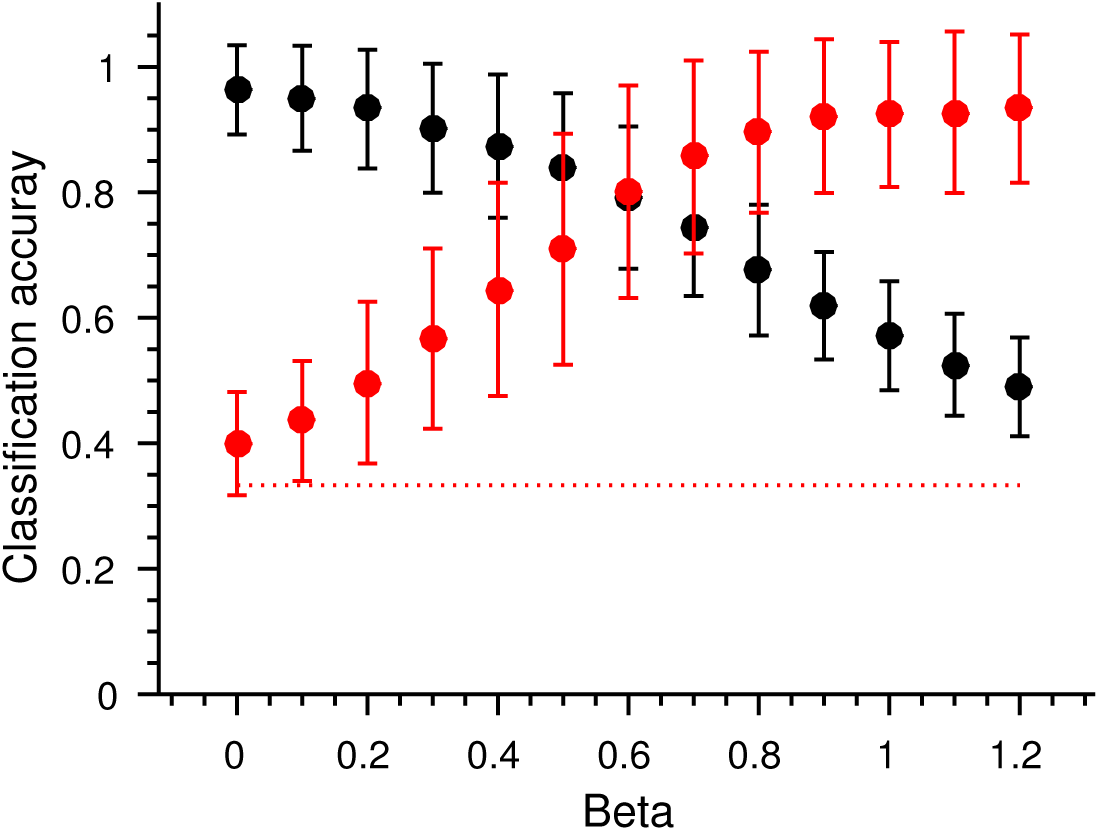
Linear classification accuracy of item identity (black) and position (red) as a function of additive interference (as represented by the *β* parameter, Eq. 2). The red dotted line shows chance level classification accuracy. Error bars depict SEM based on 1,000 simulations of the interference process with fixed parameter values.

#### 4.1.2 Positional pattern similarity decreases as a function of lag

Interference between item representations results in a change in pattern similarity across sequence positions. Specifically, between-position pattern similarity decreases as the distance between positions (lag) increases. In other words, pattern similarity is significantly higher across items that shared the same temporal position information than between items that are 1 or more positions apart (Figure 13). For the purposes of creating more positions the following plot (Figure 13) displays data generated exactly as above but with 5-item sequences instead of three.

**Figure 13:**
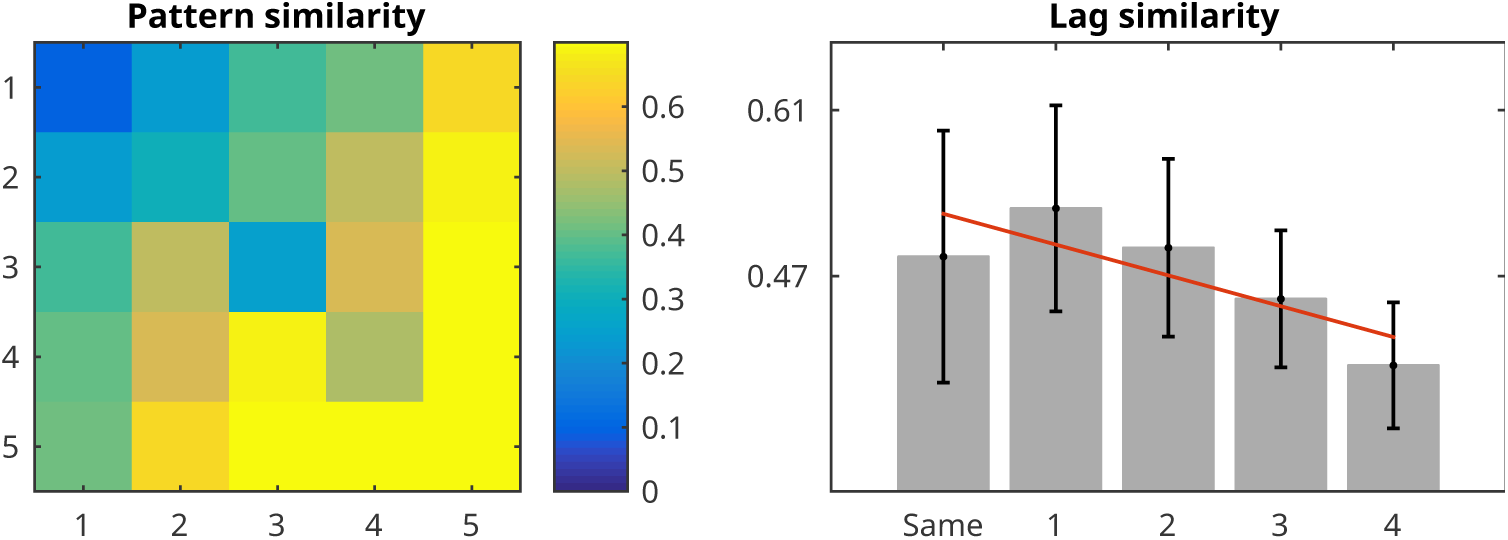
Similarity matrix on the left shows average positional pattern similarity, as measured by Pearson’s *ρ*, based on additive interference with *β* = 0.8. Plot on the right visualises this similarity as a function of positional lag. The red line depicts a statistically significant negative slope over positional lag (*p <* 0.05).

Such an effect of positional pattern similarity has be observed in a number of animal and human studies (Devito & Eichenbaum, 2011; Fortin, Agster, & Eichenbaum, 2002; Hsieh, Gruber, Jenkins, & Ranganath, 2014; Hsieh & Ranganath, 2015) and interpreted as a signature of positional code. The size of the lag effect can be measured as the magnitude of the negative slope over lag values as depicted on Figure 13 (right). Since positional effects are here solely caused by the interference mechanism it follows that the size of the lag effect correlates with the *β* parameter, which determines the extent of residual activity from the previous item. Figure 14 shows how the change in the lag effect as a function of additive interference.

**Figure 14:**
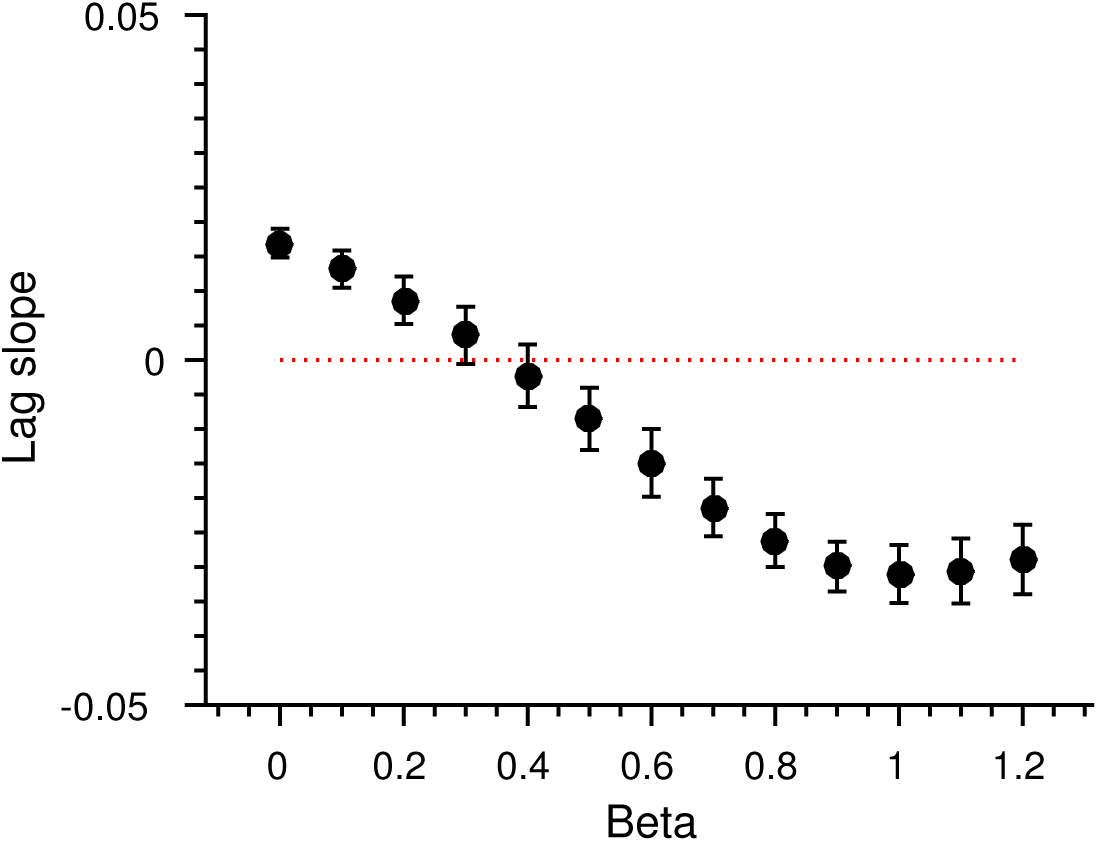
The size of the positional lag effect as a function of additive interference (*β*). Error bars depict SEM based on 1,000 simulations of the interference process with fixed parameter values.

### 4.2 Proportional interference

Pure additive interference is implausible since it presumes unlimited growth of the response in the brain region. We can cap the total response in the brain region (Σ**y**_*p*_) by normalising the response pattern every time a new item is presented. The easiest way to do this is to change the role *β* from the amount of residual activity to the *proportion* of residual activity. This requires a single change to the interference mechanism (Eq. 1) so that now we also weigh the current item representation r_*i*_, but with 1 *− π*:

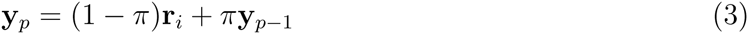

Although the mixing coefficient *π* is here calculated exactly as before (Eq. 2: *π_p_* = *π*_0_*β^p−^*^1^) its meaning has changed. Whereas previously *β* represented the amount of interference from the previously presented item, now *β* determines the *proportion* of the previous item pattern y_*p−*1_ in the current item pattern y_*p*_. If we set *β* = 0.2, the representation of a four-item sequence *A, B, C, D* would evolve as follows:

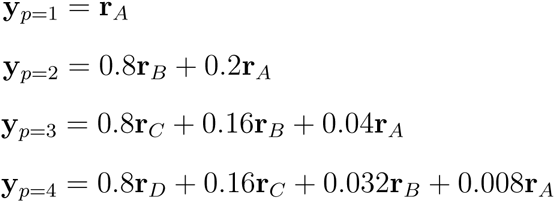

Though the mechanism of interference here is the same as in the previous simulation, we no longer allow the response of the brain region (Σ**y**_*p*_) to grow as the sequence proceeds. In other words, we have eliminated any univariate signal correlated with position. Consequently, linear decoding of response patterns based on position is not significantly different from chance any more (Figure 15A, red line). However, the positional lag effect remains since it is based on pattern similarity (as measured by Pearson’s *ρ*) which is insensitive to class means (Figure 15B).

**Figure 15:**
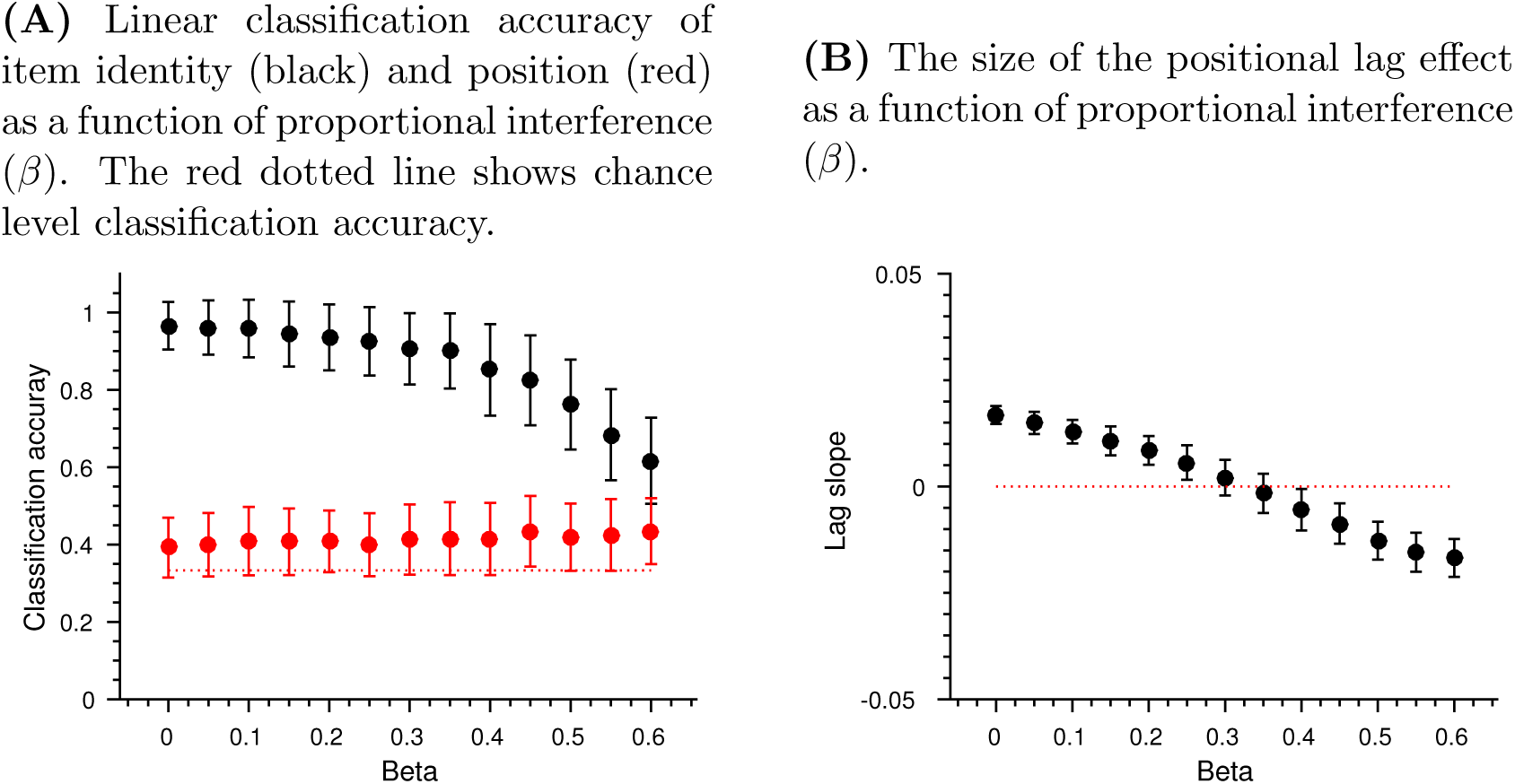
Classification accuracy and positional similarity as simulated by the proportional interference mechanism. Error bars depict SEM based on 1,000 simulations of the interference process. Notice that *β* values on the x-axis have been approximately halved since the parameter now indicated the proportion of residual activity.

In sum, de-meaning the neural response patterns only subtracts univariate effects of between-item interference. Pattern similarity effects of interference, such as the positional lag effect, still remain. It follows therefore that the positional lag effect alone is not a sufficient evidence for a neural positional code and additional statistical tests, such as classification analysis, are required.

### 4.3 Summary of item interference effects

Even in the absence of any true positional code, if the encoding of item information is based on overlaying item representations in a non-additive fashion this can potentially masquerade as a positional code. Depending on the magnitude of interference both position decoding and positional lag effects can be successfully simulated. Positional decoding is possible when residual activity from previous items is not capped and the brain region’s mean response grows with sequence position. When the activity patterns are normalised so that the mean response stays the same then only the positional lag effect remains.

## 5 Other sources of interference

The mechanism of interference, as described above in the context of item codes, can be similarly applied to other variables of the experimental design. In fact, as outlined in the *Introduction*, any fixed parameter of the experimental design is collinear with positional effects. Next we briefly discuss how position-like codes emerge as a result of interference between task phases and as a result of temporally convolved measurement.

### 5.1 Interference between task phases

One of the most common tasks used in studying sequence representation is the *serial recall task* (Figure 16). In the serial recall task presentation of a sequence of items is usually followed by a response phase requiring the participant to recall the sequence. Importantly, the temporal order between task phases themselves is always fixed: recall must necessarily follow presentation, rest always occurs between the trials etc. As a result, the positional structure of the presented sequence in the task is collinear with the structure of the task itself. For example, in the serial recall task the last item in the sequence is always followed by the recall phase. Similarly, the first item in the sequence is always preceded by recall on the previous trial. As a result we can reliably predict the position of an item in the sequence based on its adjacency to different task phases.

**Figure 16:**
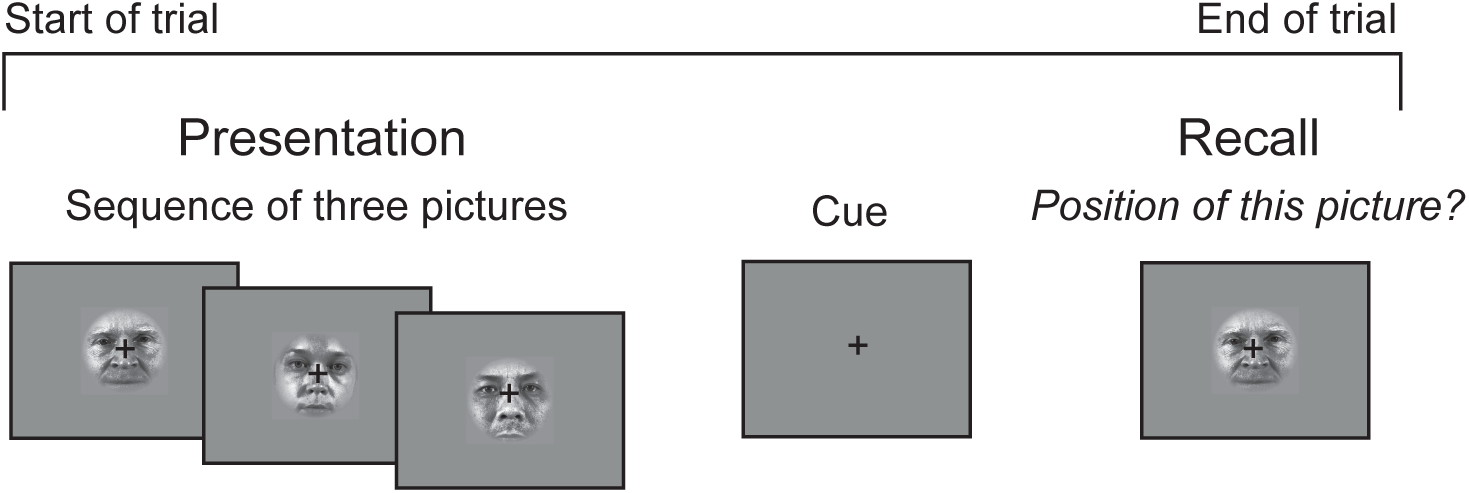
A serial recall task based on Kalm and Norris (2016).

We can model interference between task phases by simulating a response of 20 voxels as above, but during a single trial of a serial recall task. The task has two phases occurring in fixed order: *presentation* and *recall* (Figure 16). We assume that some voxels in the brain region are selective for the presentation and some for the recall phase. This selectivity can be described as voxels’ likelihood to respond given a task phase. If there is no interference between task phases the response of phase-selective voxels is independent at any stage of the task: the previous phase of the task does not alter the voxels’ activity at current stage (Figure 17A). However, if we implement additive interference as described above then the extent of the response of phase-selective voxels becomes collinear with item position in the sequence (Figure 17B). Importantly, no item codes are necessary here, just sensitivity to task phases suffices. Due to interference we can now linearly separate the response patterns in terms of their sequence *position* because the total response changes as a function of task phase (Figure 17B). In every other aspect the mechanism is the same as described in *Positional code from interference* above.

**Figure 17:**
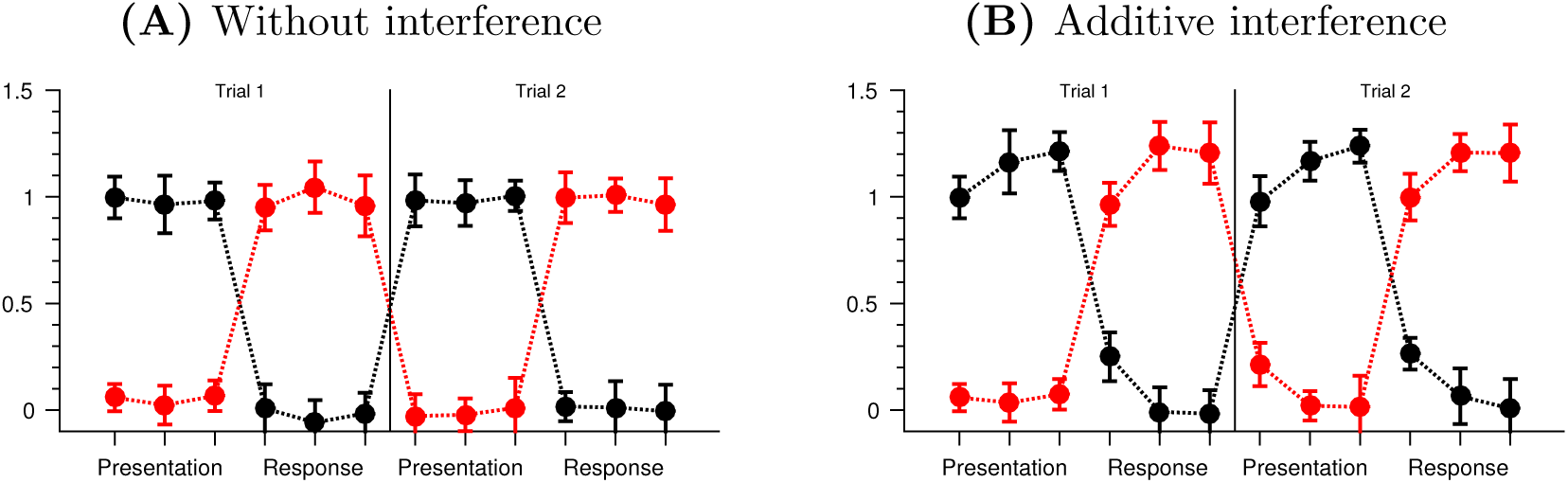
The average simulated activity of two sets of voxels, each sensitive either to the presentation or recall phase of the task. In this hypothetical task a presentation of three items in a sequence is followed by recall of three items.

Neurons’ or voxels’ sensitivity to a specific task phase is common, since in most experiments perceptual information is impossible to balance across task phases. For example, the presentation phase in serial recall task commonly uses a different stimulus modality (visual or auditory) than the following recall phase (manual or verbal recall, see Table 1). As a result, large patches of the cortex are only engaged during a specific phase of the task creating conditions described above.

### 5.2 Interference from measurement: functional MRI

So far we have described interference mechanisms arising between neural representations. However, equally importantly, interference between representations can result from noisy measurement. Similarly to representational interference, this can lead to positional effects which are spurious.

Functional MRI measures neural activity by detecting changes in the concentration of oxyhemoglobin and deoxyhemoglobin in neural tissue (BOLD signal). The relationship between a neural event and the corresponding BOLD signal can be described by a haemodynamic response function (HRF). Importantly, the HRF is non-linear and spread out over several seconds (Figure 18A), meaning that the BOLD signal corresponding to temporally adjacent events, such as items in a sequence or task phases, will always contain a response elicited by events preceding the event of interest (Figure 18B). This creates conditions similar to between-item and task phase interference described above – only this time there is no need for cognitive or representational interference. The temporal overlap in the BOLD signal will result in interference between measured item or phase representations even if the neural representations are independent of each other.

**Figure 18:**
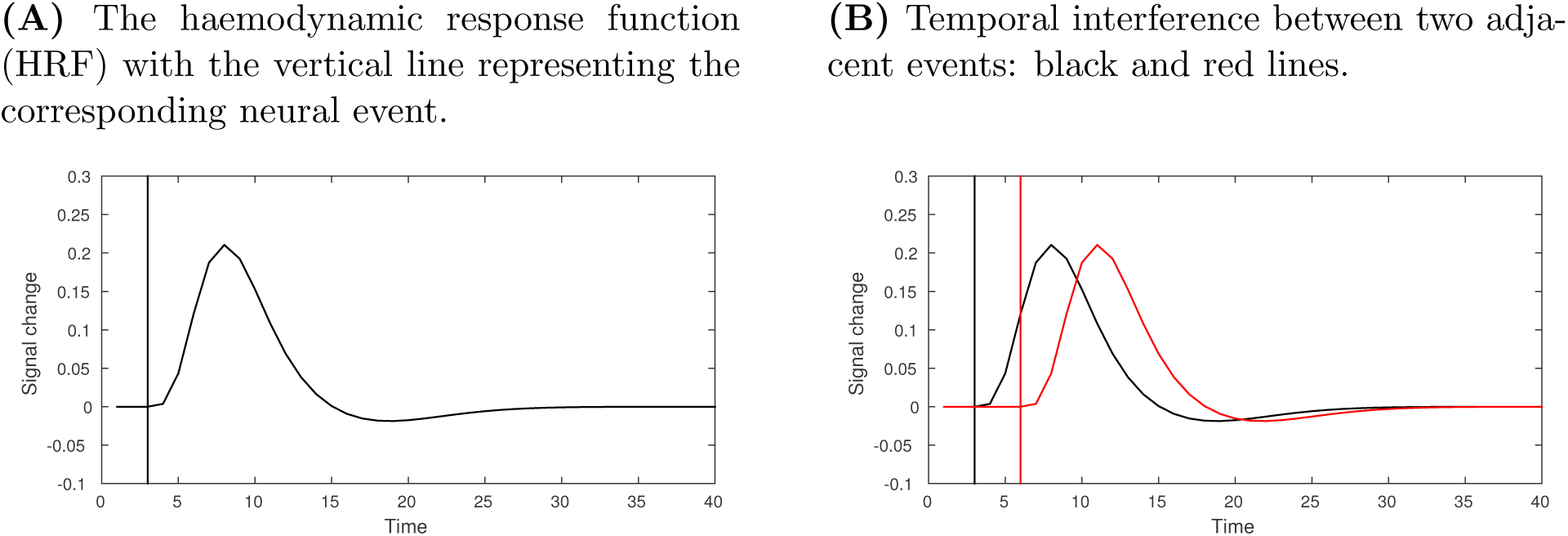
Temporal interference in fMRI

## 6 Discussion

A major methodological barrier to studying the neural representation of positional code is that in any sequence processing task items in different position necessarily differ on other dimensions too. In this paper we used simulations and experimental data to show how both position-collinear experimental variables, noisy measurement, and interference between sequence items can lead to positional read-out in the absence of a dedicated positional code. This raises two important questions: (1) is it important to distinguish between a positional read-out and a ’true’ positional code; and (2) what steps can be taken to delineate those in experimental data.

### 6.1 Positional read-out vs. dedicated positional code

Most models of neural sequence representation assume the existence of a dedicated positional code in the brain (Howard & Kahana, 2002; Henson & Burgess, 1997; Burgess & Hitch, 2006; Brown, Preece, & Hulme, 2000). However, since several cognitive processes (e.g. memory load, sensory adaptation) are collinear with any positional signal, a question arises whether those collinear processes could be used as a positional signal not just by the experimenter but also by the brain. We argue that a positional read-out from either simple position-collinear processes or between-item interference is not sufficient to support the storage and recall of a sequence.

### 6.1.1 Positional read-out from collinear processes is not sufficient for recalling a sequence

Recalling items in a sequence requires reinstating their order during recall. This problem is solved by positional models of sequence processing by associating each sequence item with its position during encoding and retrieving the order of items during recall by reinstating each positional code, which then cues the associated item (Figure 1B, e.g.: Burgess & Hitch, 2006; Howard & Kahana, 2002; Brown et al., 2000; Lee & Estes, 1981). However, it is hard to see how processes such as memory load or sensory adaptation could be used to cue associated items. Although experimenters can decode item position in a sequence based on memory load or sensory adaptation it is hard to see how ”cortex as receiver” can use those neural signals to represent position and guide behaviour. For example, in case of using memory load (or any monotonically changing signal) as a positional code to cue associated items would require first to reinstate such ’load’ to cue the corresponding item. However, such interpretation of ’memory load’, which can be reinstated independently of the amount of items in memory, loses its original meaning and becomes a clumsy re-interpretation of a dedicated positional code. For this reason any *effect* of sequence processing, such as memory load or sensory adaptation, cannot be inverted into *cause* that would enable to associate items into sequence.

### 6.1.2 Positional read-out from between-item interference is not sufficient for re-calling a sequence

We showed that interference between item representations can potentially masquerade as a positional code. This is because noisiness of the item representations changes monotonically over sequence positions as a result of interference. The change in the noise profile can therefore be used by the experimenter to reliably predict the position of the item in the sequence. However, as with simple position-collinear processes, it is hard to see how changes in the signal-to-noise ratio can be used by the brain to guide sequence recall. The main consequence of interference via overlaying item representations is that the later items in the sequence are noisier than the early ones. This contradicts the well-established recency effect in sequence recall, where last items in the sequence are more accurately recalled than the ones in the middle (see Hurlstone, Hitch, & Baddeley, 2014, for a review). Secondly, using the noisiness of item representations as a positional code to cue associated items conflates the cause and effect relationship in sequence processing, as discussed above. The noisiness of the items would need to be reinstated independently of items themselves, thus necessitating the recoding of the noise levels into a dedicated positional signal.

### 6.2 Methods to dissociate between positional read-out and dedicated positional code

It is not possible to devise a task where the positional signal is orthogonal to other experimental variables since cognitive processes collinear to the positional code will always arise whenever stimuli are presented in sequence. However, the vast majority of previous studies on the positional code (Table 1) do not acknowledge the possibility of the ’contamination’ of the positional code or take any measures to control for it.

Two assumptions are required to distinguish between a positional read-out and a ’true’ positional code. First, position-collinear processes like memory load or sensory adaptation will uniformly affect all neural units engaged in encoding the item representations. This assumption is relatively uncontroversial if we presume that such processes are the result (and not source) of sequence processing. Second, we need to assume that a dedicated positional code is reflected in the position-sensitivity within a population of neural units. In other words, units respond selectively to sequence positions based on some tuning function. Under such conditions simple de-meaning (e.g z-scoring) of the neural data with respect to experimental condition (item position) will eliminate any univariate signal from the data including any univariate positional read-outs (see *Eliminating uniform signal by de-meaning*).

However, we also showed that between-item interference can result in pattern similarity effects which masquerade as positional signal in the form of the lag effect (see *Positional pattern similarity decreases as a function of lag* and *Proportional interference*). Effects of pattern similarity are independent of signal amplitude and hence invariant to de-meaning. As a result, the effect of positional lag which has been used in several previous studies of positional code (Devito & Eichenbaum, 2011; Fortin et al., 2002; Hsieh et al., 2014; Hsieh & Ranganath, 2015) cannot be taken as a proof of neural positional code without ruling out between-item interference first. We show that this can be achieved by using linear classification analyses on the de-meaned neural responses.

Besides cognitive interference– such as based on overlaying item representations – positional read-out can result in noisy measurement, such as the temporal interference inherent in fMRI. In other words, any fMRI signal pertaining to successively presented sequence items will include a positional signal based on measurement error, even if we assume no interference between the neural representations of items themselves. As a result, the positional lag effect alone should never be used in fMRI studies as an indicator of neural positional representation. In fMRI studies sequentially presented stimuli will always be collinearly dependent on each other because of the inherent temporal lag in the BOLD signal. As a solution, whole-sequence data can be used to extract positional information using the representational similarity analysis (Kriegeskorte, Mur, & Bandettini, 2008; Kalm & Norris, 2014).

### 6.3 Conclusions

In this paper we have explored two types of processes that could enable an experimenter to read out a positional ’code’ in the absence of a dedicated positional code. First, we show that with any sequence processing task there are experimental variables collinear with the positional signal (e.g. time, memory load, etc.) which can serve as a positional code. Second, we show how interference between item representations, task phases, and measurement modalities can also lead to a similar positional read-outs.

We argue that it is important to distinguish between a positional read-out and a dedicated positional code, since only the latter has been shown to be compatible with experimental data. Furthermore, we argue that such collinear processes which enable positional read-out are the result of sequence representation not cause, and hence would not be able to even theoretically support sequence retrieval. Finally, we suggest practical steps in data analysis to distinguish between a positional read-out and a code. Furthermore, this paper shows that many results from behavioural and neural experiments studying the positional code must be treated with caution.

The MATLAB/Octave code for the simulated data and plots is freely available at http://imaging.mrc-cbu.cam.ac.uk/imaging/KristjanKalm/ovs/.

**Table 1:** Studies of neural representation of positional code

| First author | Year | Stimuli | Task | Subject | Measurement |
| --- | --- | --- | --- | --- | --- |
| Amiez | 2007 | visual | manual | human | fMRI |
| Averbeck | 2003 | motor | motor | monkey | electrophysiology |
| Averbeck | 2006 | visual | saccade | monkey | electrophysiology |
| Averbeck | 2007 | visual | saccade | monkey | electrophysiology |
| Barone | 1989 | visual | manual | monkey | electrophysiology |
| Berdyeva | 2010 | visual | saccade | monkey | electrophysiology |
| Berdyeva | 2011 | visual | motor | monkey | electrophysiology |
| Carpenter | 1999 | visual | motor | monkey | electrophysiology |
| Crowe | 2014 | manual | manual | monkey | electrophysiology |
| DuBrow | 2014 | visual | manual | human | fMRI |
| DuBrow | 2016 | visual | manual | human | fMRI |
| Fujii | 2005 | visual | saccade | monkey | electrophysiology |
| Gelfand | 2003 | auditory | manual | human | fMRI |
| Nieder | 2006 | visual | manual | monkey | electrophysiology |
| Ginther | 2011 | odour | motor | rodent | electrophysiology |
| Heusser | 2016 | visual | manual | human | MEG |
| Hsieh | 2014 | visual | manual | human | fMRI |
| Hsieh | 2015 | visual | manual | human | fMRI |
| Hyde | 2012 | in vitro |  | rodent | electrophysiology |
| Inoue | 2006 | visual | manual | monkey | electrophysiology |
| Isoda | 2004 | visual | saccade | monkey | electrophysiology |
| Kalm | 2014 | auditory | auditory | human | fMRI |
| Kalm | 2016 | auditory | visual | human | fMRI |
| Kraus | 2013 | motor | motor | rodent | electrophysiology |
| Lehn | 2009 | visual | manual | human | fMRI |
| MacDonald | 2013 | odour | motor | rodent | electrophysiology |
| MacDonald | 2011 | odour | motor | rodent | electrophysiology |
| Mankin | 2012 | spatial | motor | rodent | electrophysiology |
| Manns | 2007 | odour | motor | rodent | electrophysiology |
| Manns | 2007 | odour | motor | rodent | electrophysiology |
| Merchant | 2013 | auditory | manual | monkey | electrophysiology |
| Nakajima | 2009 | manual | manual | monkey | electrophysiology |
| Naya | 2011 | visual | manual | monkey | electrophysiology |
| Nieder | 2012 | visual | manual | monkey | electrophysiology |
| Ninokura | 2004 | visual | manual | monkey | electrophysiology |
| Pastalkova | 2008 | motor | motor | rodent | electrophysiology |
| Petrides | 1991 | visual | manual | monkey | lesion |
| Rangel | 2014 | motor | motor | rodent | electrophysiology |

